# Electrical and chemical synapses share similar organizational principle

**DOI:** 10.64898/2026.05.19.726377

**Authors:** Hannah Hoff, Sundas Ijaz, Fabio A. Echeverry, Stephan Tetenborg, Ya-Ping Lin, John O’Brien, Vytas K. Verselis, Alberto E. Pereda

## Abstract

Electrical transmission is mediated by intercellular channels that cluster into structures known as ‘gap junctions’ (GJ). In vertebrates, GJ channels are encoded by the gene family of connexin (Cx) proteins that assemble as hexamers, termed hemichannels, in the pre- and postsynaptic membranes, and that subsequently dock to form GJ channels. Auditory contacts on the fish Mauthner cells serve as model to study the properties and organization of vertebrate electrical synapses. Electrical transmission at these synapses is mediated by multiple co-existing GJs at which the presence of intercellular channels is regulated by a molecular scaffold. Zebrafish contain four homologs of the neuronal Cx36: Cx35.5 and Cx35.1 (*gjd2a* and *b*, respectively), and Cx34.1 and Cx34.7 (*gjd1a* and *b*). Cx mutations suggested that GJs are formed by heterotypic channels made of presynaptic Cx35.5 and postsynaptic Cx34.1. Using transgenic fish in which Cxs were tagged, we found that a second Cx, Cx34.7, is present together with Cx34.1 on the postsynaptic side at some but not all GJs at these terminals. When exogenously expressed, both Cx34.1 and Cx34.7 formed heterotypic functional channels with Cx35.5, each with substantially different voltage-dependent properties, indicating they can serve differential functions. However, we previously demonstrated that electrical transmission is lost in Cx34.1 but not Cx34.7 null mutants, suggesting that Cx34.7 cannot compensate for the loss of Cx34, despite the intrinsic ability of Cx34.1 and Cx34.7 to create functional channels. The findings reveal an unanticipated functional organization in the electrical synapse, where Cx34.1 is obligatory and Cx34.7 accessory, roles that appear to be defined by the postsynaptic molecular scaffold, with two postsynaptic Cxs possibly assembling under specific functional contexts. Thus, our results indicate that electrical synapses share an organizational motif with chemical synapses, akin to how they combine postsynaptic receptor types to modify synaptic function.

## INTRODUCTION

Communication between neurons takes place at functional contacts known as ‘synapses’. Chemically-mediated synapses are recognized as structurally and molecularly complex (1). Although presynaptic specializations make transmitter release and its regulation possible, an intricate molecular postsynaptic machinery ultimately shapes transmission by combining functionally different neurotransmitter receptors, thus bringing diversity to chemical synaptic communication (2–5). Electrical synapses are, on the other hand, considered simpler and mediated by intercellular channels which cluster into a structure, or ‘plaque’, known as a ‘gap junction’ (GJ), that has been presumed to function autonomously in mediating the direct spread of electrical currents and small metabolites between neurons (6). The communicating intercellular GJ channel is formed by the apposition of two hemichannels, each contributed by one of the communicating cells. However, emerging evidence indicates that electrical synapses are molecularly and structurally complex and share forms of organization with their chemical counterparts (7–13). Accordingly, recent proteomic analyses in zebrafish and mammalian systems (12, 13) have revealed an abundance of scaffolding proteins and additional synaptic machinery associated with neuronal GJs comparable to that of the postsynaptic density (PSD) associated with glutamatergic synapses (14).

Auditory afferents terminating as single ‘Large myelinated Club endings’ (CEs) on the lateral dendrite of the Mauthner cell (M-cell; a pair of colossal reticulospinal neurons that mediate sensory-evoked tail-flip escapes responses; (15)) contain both GJs and specializations for chemical transmission (16, 17). Because of their experimental access, these identifiable auditory contacts have served as a valuable model to explore the properties of vertebrate electrical synapses (17–21). Electrical transmission at these terminals is mediated by not one GJ, but multiple (∼35) co-existing GJs that collectively occupy ∼80% of the contact area (22). GJ channels at these terminals are formed by connexins (Cxs) Cx35 and Cx34, teleost homologs of the mammalian Cx36, with their presence shown to be under the control of associated scaffolding proteins (23–25). Electrical transmission at CEs is dynamically regulated and undergoes activity-dependent potentiation (19–21). These plastic properties result from a close functional relationship between GJs and the co-existing glutamatergic specializations for the induction of electrical synaptic plasticity (26), which was later reported to occur in mammals as well (27–29). Thus, the properties and molecular organization of electrical synapses at CEs has shed light onto the molecular organization of vertebrate electrical transmission, in general.

Leveraging the experimental accessibility of CEs in zebrafish with the capacity for genetic manipulation, we investigated the Cx organization of GJs in these terminals. The zebrafish brain contains four neuronal connexins, Cx35.5 and Cx35.1 (genes *gjd2a* and *gjd2b*) and Cx34.1 and Cx34.7 (genes *gjd1a* and *gjd1b*), with distinct localization patterns (30). Mutations of Cx genes combined with chimeric analyses (30) suggested that GJs at CE contacts are formed by heterotypic channels made of presynaptic Cx35.5 hemichannels and postsynaptic Cx34.1 hemichannels (25, 30). Using transgenic fish in which Cxs were tagged with fluorescent proteins, we confirmed that Cx35.5 is the presynaptic connexin. However, we find that Cx34.7 is present postsynaptically in combination with Cx34.1 at some but not all the GJs within these terminals. This finding indicates an additional asymmetry in the composition of the GJ plaques at CE terminals where, in contrast to the presynaptic side, the hemiplaques on the postsynaptic side express two Cx orthologs. Because Cx34.7 was shown unable to compensate for the loss of transmission in Cx34.1 null mutants, the finding reveals a novel functional organization for GJ channels, where Cx34.1 plays an obligatory role and Cx34.7 plays an accessory role, potentially assembling with Cx34.1 in a heteromeric fashion. Our results also show that both Cx34.1 and Cx34.7, when exogenously expressed, form channels with Cx35.5, although with different biophysical properties. This finding indicates that this Cx configuration does not result from an intrinsic inability of Cx34.7 to form functional channels with Cx35.5 but, rather, likely to be instructed by the molecular machinery of the postsynaptic site of the CE electrical synapse. Because of its biophysical properties that are distinct from Cx34.1, Cx34.7 is likely to operate in specific functional contexts imparting different functional properties to GJs of the CE electrical synapse.

## RESULTS

The Large Myelinated Club endings (CEs) are an identifiable group of auditory synaptic contacts on the Mauthner (M-) cells of fish (16), including larval zebrafish (ZF) (30–33). Their unusually large size make them ideal for structural/functional correlations during investigations of synaptic mechanisms. At these terminals, transmission is mediated by multiple coexisting gap junctions (GJs) and specializations of glutamatergic transmission. Stimulation of these afferents evokes a ‘mixed’ synaptic response composed by an early GJ-mediated electrical component followed by small and longer lasting chemical synaptic component (Fig. 1A) (31, 34, 35). Because the multiple individual GJs collectively occupy ∼80% of the terminal surface (33), their contact areas can be easily identified in 5-6 dpf larval ZF by immunolabeling with antibodies against connexins (Cx), the GJ channel forming proteins in vertebrates. The contacts appear as large oval areas of intense staining, ∼2 µm in diameter in the distal portion of the M-cell lateral dendrite (Fig. 1B-D). Fish have been shown to contain two homologs of the mammalian neuronal Cx36: Cx34 (*gjd1*) and Cx35 (*gjd2*). Using Freeze fracture immunolabeling (FRIL) double replica, Cx35 was found to be limited to presynaptic hemiplaques and Cx34 to postsynaptic (dendritic) hemiplaques at CEs in goldfish (9) and larval ZF (11), as well other hindbrain synapses (10). Due to the genome duplication in teleosts, ZF contains four neuronal connexins, Cx35.5 and Cx35.1 (genes *gjd2a* and *gjd2b*) and Cx34.1 and Cx34.7 (genes *gjd1a* and *gjd1b*), each with distinct distribution patterns (30). Chimeric analyses of mutations of Cx genes confirmed the presynaptic distribution of Cx35 and Cx34, and suggested that GJs at CEs are formed by heterotypic channels made of presynaptic Cx35.5 hemichannels and postsynaptic Cx34.1 hemichannels (25, 30). These results are consistent with the lack of immunolabeling for Cx35.5 and Cx34.1 (see Fig. 2 in Ref. (25)) in *gjd2a/Cx35.5-/-* and *gjd1a/Cx34.1-/-* mutant ZF and an accompanying lack of a detectable early electrical component in the mixed synaptic response observed in WT; only the delayed chemical component remained (See Fig. 3C–E in Ref. (25)). In contrast, electrical transmission was unaffected in *gjd2b/Cx35.1-/-* and *gjd1b/Cx34.7-/-* mutant zebrafish (see Figure 3E; figure supplement 1A-B in Ref.(25)), a finding supported by immunolabeling in these fish (see Fig. 2; figure supplement 1G-L in Ref. (25)).

**Figure 1.**
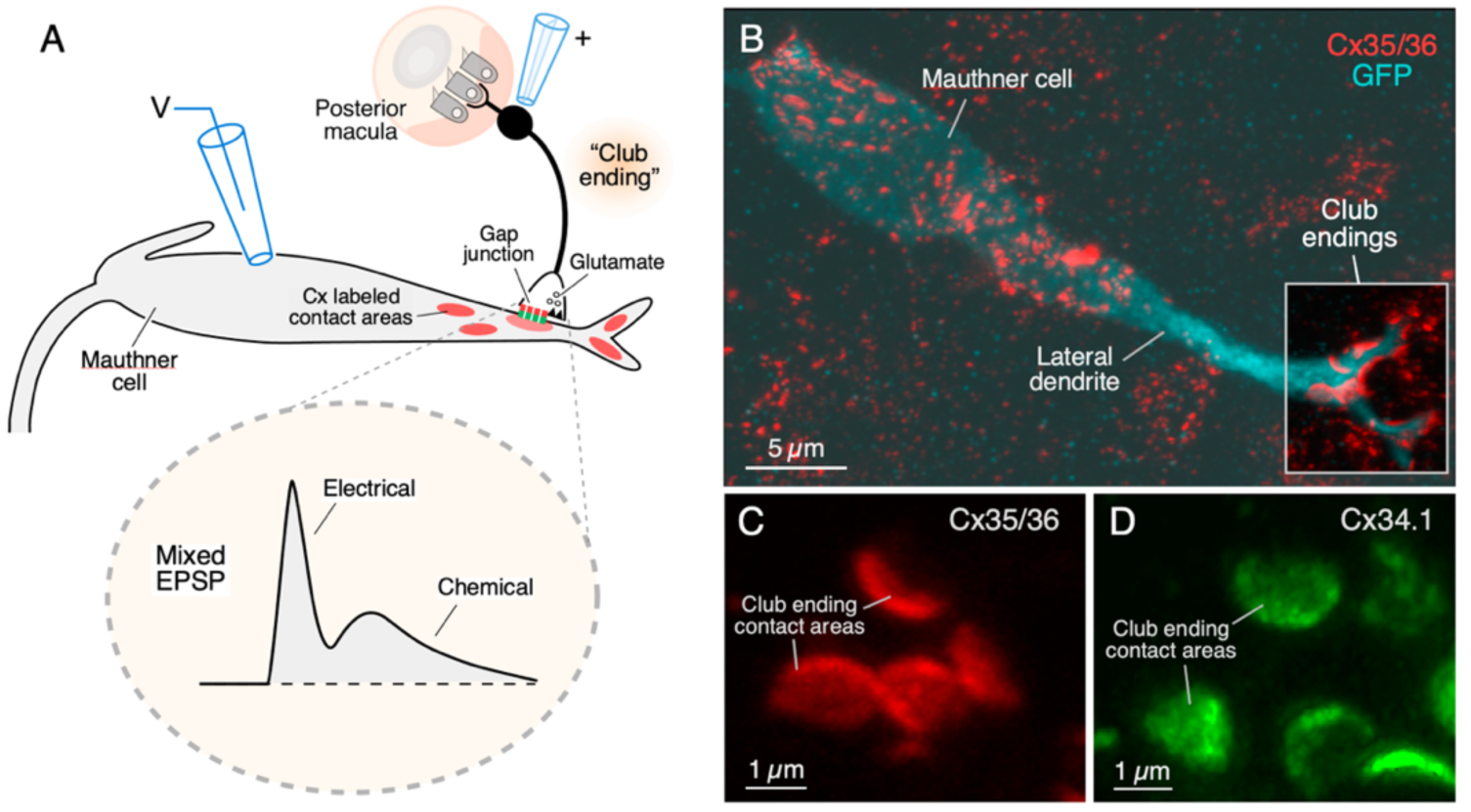
Club endings are identifiable synapses on the Mauthner cells of larval zebrafish. (A) The cartoon illustrates the auditory afferents that terminate as single Club endings on the distal portion of the lateral dendrite of the Mauthner (M-) cell of zebrafish larvae, each containing gap junctions (gap junction) and specializations for chemical transmission (glutamate). Synaptic contact areas labeled with connexin antibody (see B, C, D) are represented in red. The diagram also illustrates the experimental paradigm used to examine synaptic transmission. Extracellular stimulation (+) of auditory afferents at the posterior macula evokes ‘mixed’ synaptic response (Mixed EPSP) in the M-cell (V), composed of an early electrical component mediated by the gap junctions (Electrical) followed a delayed glutamatergic response (Chemical). Modified from Lasseigne et al., Elife, 2021(25). (B) Confocal image with anti-GFP (cyan) and anti-Cx35/36 (red), which labels both Cx35.5 and Cx35.1, showing the soma and a long stretch of the lateral dendrite of the M-cell (projection of 42 confocal z-sections at 0.40 µm z-step size). Box: the Club endings are segregated to the distal portion of the lateral dendrite (the image delimited by the box was cropped from and placed on the same region of the lighter image). (C-D) Contact areas of individual CEs labeled with anti-Cx35/36 (C, red; projection of 4 sections at 0.40 µm z-step size) and anti-Cx34.1 (D, green; projection of 20 sections at 0.19 µm z-step size).

**Figure 2.**
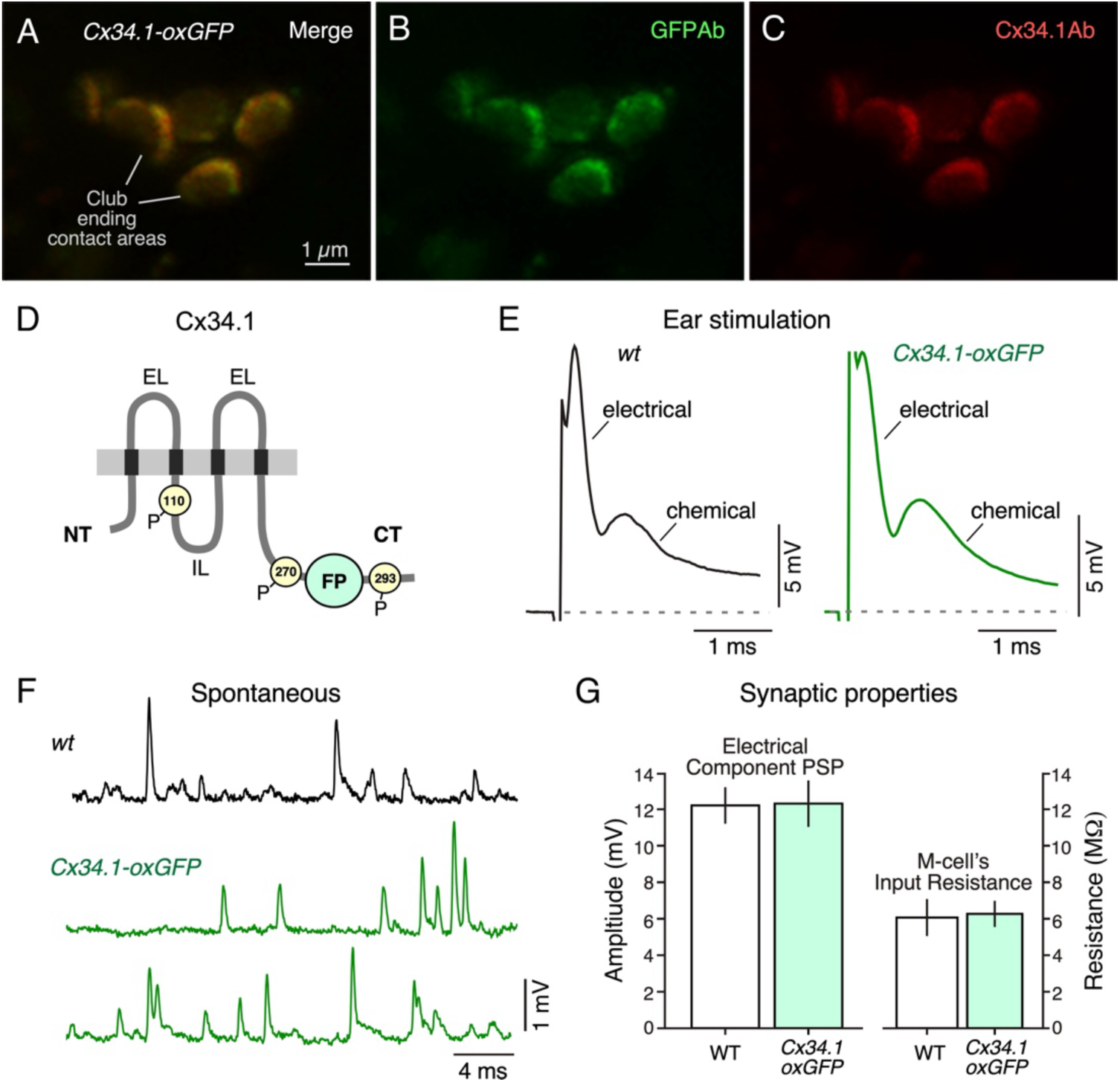
Generation of transgenic zebrafish at which connexins are tagged. (A) Club ending contact areas labeled with anti-GFP (green) and anti-Cx34.1 (red) in a Cx34.1-oxGFP 5 dpf transgenic zebrafish (projection of 13 confocal z-sections at 0.40 µm z-step size). (B) Club ending contact areas labeled with anti-GFP. (C) Club ending contact areas labeled with anti-Cx34.1. (D) Transgene design. Cx34.1 membrane topology: NT: amino terminus; CT: carboxy terminus; IL: intracellular loop; EL: extracellular loops; P: phosphorylation sites. The fluorescent protein (FP) coding region was inserted into the CT of the connexin genes at a site (19-21 aa before end of the CT between phosphorylation sites at aa 270 and 293), which was previously found to preserve regulation of coupling by connexin phosphorylation and as well as the CT PDZ-domain interaction motif. (E) Electrophysiological recordings in these animals exhibit normal transmission. (E) Mixed synaptic response in the M-cell evoked by stimulation of CE afferents near the posterior macula of the ear for a wild type (WT, black trace) and a Cx34.1-oxGFP (green trace) 5 dpf ZF (recordings represent an average of at least 10 synaptic responses). (F) Spontaneous synaptic responses obtained in a WT (top) and Cx34.1-oxGFP ZF (bottom). (G) Quantification of synaptic and electrical properties. Left: amplitude of electrical component. WT: 12.2 ± 1.0; Cx34.1-oxGFP: 12.3 ± 1.2 (Mean ± SEM mV); Right: Input resistance of M-cells. WT: 6.1 ± 1.0; Cx34.1-oxGFP 6.3 ± 0.7 (Mean ± SEM MOhm); n=5 fish, p=0.75 and 1 (Mann-Whitney), respectively.

**Figure 3.**
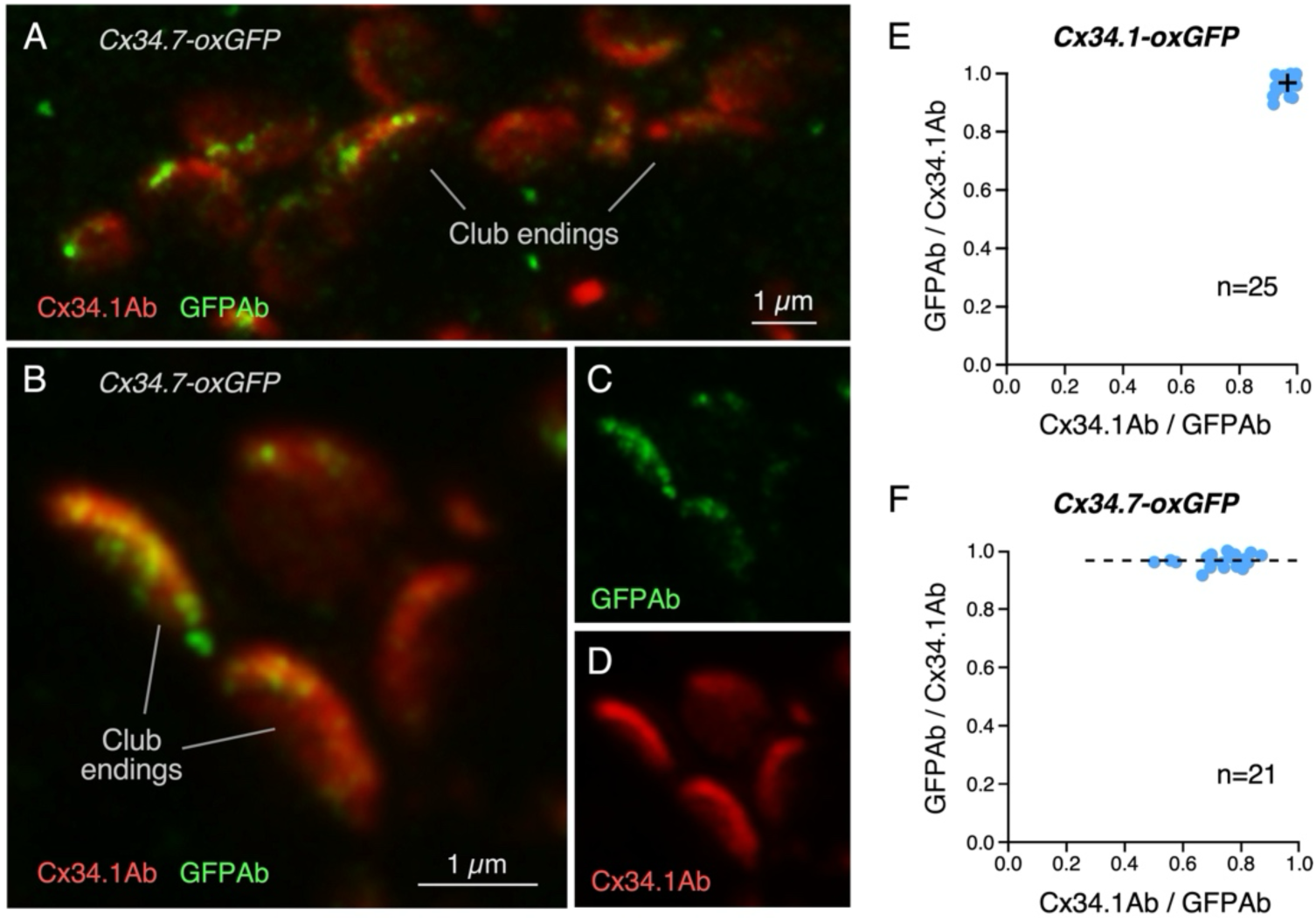
Transgenic zebrafish show the presence of Cx34.7 at Club endings. (A) CE synaptic contact areas labeled with anti-Cx34.1 and anti-GFP on a stretch of the lateral dendrite of a Cx34.7-oxGFP ZF (projection of 4-26 sections at 0.40 µm z-step size). (B) Magnification of contact areas labeled with anti-Cx35.5 and anti-GFP showing different amounts of Cx34.7 between neighboring CEs (projection of 4-20 sections at 0.40 µm z-step size). (C) Labeling with anti-GFP. (D) Labeling with anti-Cx34.1. (E) Graph showing colocalization of GFP and Cx34.1 fluorescence at individual CEs in Cx34.1-oxGFP ZF determined by the Mander’s Coefficient: GFPAb/Cx34.1Ab 0.968 ± 0.030 (x axis); Cx34.1Ab/GFPAb 0.965 ± 0.025 (y axis), n= 25 CEs from 14 fish. Cross mark indicates the average value. (F) Colocalization of GFP and Cx34.1 fluorescence at individual CEs in Cx34.7-oxGFP fish determined by the Mander’s Coefficient: GFPAb/Cx34.1Ab 0.966 ± 0.022 (x axis); Cx34.1Ab/GFPAb 0.729 ± 0.094 (y axis), n= 20 CEs from 18 fish.

To further investigate the distribution of these Cxs in the M-cell system we generated transgenic ZF in which the Cxs are tagged with fluorescent proteins (see Methods). The fluorescent protein coding regions were inserted into the CT of the connexin genes at a site, 19-21 aa before end of the CT (see Fig. 2D) that was previously found to avoid disruption of the regulation of coupling by Cx phosphorylation and to prevent block of the C-terminal PDZ-domain interaction motif (36). For this purpose, Cx34.1, the postsynaptic Cx, was tagged with oxGFP. As proof of the success of the transgenesis method: i) the transgene was expressed at the expected sites, and ii) electrical transmission at these sites was preserved. That is, the CE terminals were easily identified with an anti-GFP antibody and this labeling overlapped with anti-Cx34.1 labeling (Fig. 2A-C), which recognizes both endogenous and transgene Cxs. The mixed synaptic response evoked by stimulation of these terminals in *Cx34.1-oxGFP* animals was indistinguishable from *wt* animals (Fig. 2E). Moreover, spontaneous electrical synaptic responses including those contributed by other contacts were also consistent with those observed in *wt* animals (Fig. 2F), indicating that synaptic transmission at other sites was unaffected. We found no difference between the maximal amplitudes of the electrical component of the mixed synaptic response evoked by stimulation of CEs in *wt* and *Cx34.1-oxGFP* fish (Fig. 2G, left). Finally, the M-cell’s values of input resistance in *Cx34.1-oxGFP* fish were indistinguishable from those observed in *wt*, indicating the absence of changes in the excitability of this cell (Fig. 2G, right). An eventual reduction in the conductance of GJs at electrical synapses on the M-cell due to transgenesis would have resulted in an increase in this last value (25).

Contrasting the prediction based on immunolabeling and functional analyses of *gjd1a/Cx34.1-/- and gjd1b/Cx34.7-/-* animals, the generation of *Cx34.7-oxGFP* animals clearly revealed the presence of Cx34.7 at CEs (Fig. 3 A-C). In contrast to Cx34.1, which was distributed widely throughout all the GJs within a CE contact area, the distribution of Cx34.7 evaluated with anti-GFP was sporadic and characteristically punctate (Fig. 3A). Notably, the amount of this labeling differed among neighboring CEs (Fig. 3B-D). Interestingly, analyses of colocalization on Cx34.1 and Cx34.7 (GFP labeling) in *Cx34.7-oxGFP* fish (Fig. 3F) showed that in addition to existing at variable amounts at individual CEs, Cx34.7 always was colocalized with Cx34.1; i.e., in contrast to Cx34.1 that can be expressed alone, Cx34.7 and Cx34.1 could be co-expressed at some GJs. The co-localization analysis of Fig. 3E and F was obtained from ‘en face’ views of CE contacts in these two transgenic fish. In contrast to adult fish, ‘en face’ views in larval ZF are less likely to be found and measured because the diameters of the CE contact areas and the lateral dendrite, where the contacts are localized, are of comparable size (5, 10). The observed labeling likely represents the normal distribution of Cx34.7 because: i) using the same transgenesis method, *Cx34.1-oxGFP* fish showed a strong colocalization between GFP and Cx34.1 labeling (Fig. 3E), ii) anti-GFP labeling in *Cx34.7-oxGFP* fish revealed the presence of this Cx in the outer plexiform layer (OPL) of the retina (Supp. Fig. 1), and finally iii) this distribution was consistently observed in multiple lines of *Cx34.7-oxGFP* fish.

Our colocalization analyses suggest an intimate functional association between Cx34.7 and Cx341 in the postsynaptic hemiplaque. To investigate this possibility, we performed expansion microscopy in *Cx34.7-oxGFP* fish, a technique that allows us to explore the composition of individual GJs at CE terminals (33). Using a protocol optimized for synapses on the M-cell that yields a ∼4-fold linear expansion of CEs (see Methods and Refs (33, 38)) it is possible to observe multiple single puncta, each corresponding to a single GJ (33), defining the CE contact area (Fig. 4A-D). While most GJs were labeled only for Cx34.1, we found colocalization of Cx34.7 (GFP labeling) with Cx34.1 (Fig. 4A) and Cx35.5 (Fig. 4B; Cx35/36 antibody labeling) in multiple GJs. Figure 4C illustrates a ‘En face’ view of one on these contact areas labeled with anti-GFP and anti-Cx34.1, where GJs labeled for both Cx34.1 and Cx34.7 coexist with neighboring GJs containing only Cx34.1. In contrast, we did not observe GJs containing only Cx34.7. To further investigate the existence of these two types of GJs we performed line scans of single GJs containing either Cx34.1 alone or together with Cx34.7. The analysis was performed at puncta located at the periphery (Fig. 4D), which are more favorable for line scan analysis as a result of the characteristic concavity of the CE contact area (see Ref. (33)) and where it was more easy to observe neighboring puncta differentially labeled (Fig. 4, insets 1 and 2). Line scan of single GJs indicate the presence of labeling for either both Cx34.1 and Cx34.7 (Fig. 4E, left) or only Cx34.1 (Fig. 4E, right) at a single GJ punctum (Fig. 4C). Figure 4F shows a line scan through two neighboring puncta, one labeled for both Cx34.7 and Cx34.1 and another only for Cx34.1. The graphs of Fig. 4G and H show the average of individual line scans for double-labeled and single-labeled GJs, respectively. Thus, CE electrical synapses are formed by two types of GJs: one containing only Cx34.1, and variable amounts of second type containing both Cx34.1 and Cx34.7, with variable subregions of the terminal having GJs containing both Cxs or just Cx34.1.

**Figure 4.**
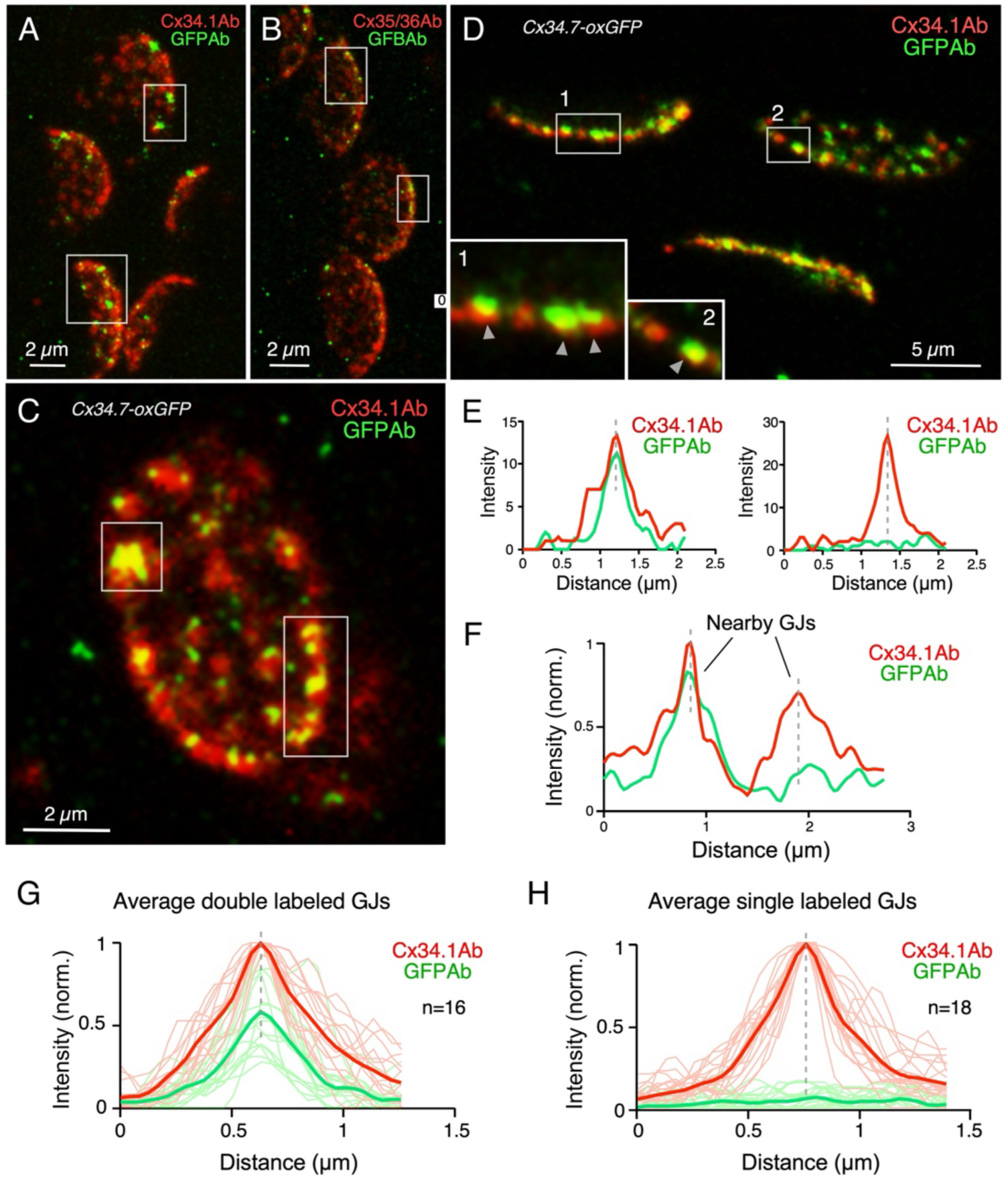
Expansion microscopy reveals the co-existence of Cx34.1 and Cx34.7 at individual gap junctions. (A) Protein-retention expansion microscopy (ProExM) with anti-Cx34.1 (red) and anti-GFP (green) increases the size of CE synaptic contact areas, enabling the visualization of subsynaptic components and revealing the distribution of Cx34.1 and Cx34.7 at individual GJs within CE contact areas in Cx34.7-oxGFP fish (projection of 30-39 sections at 0.71µm z-step size). (B) Expanded synaptic contact areas labeled with anti-GFP (green) and the Cx35/36 antibody (red), which labels both Cx35.5 and Cx35.1 (projection of 3-25 sections at.0.71 µm z-step size). (C) ‘En face’ view of an expanded contact area double-labeled for with antiCx34.1 (red) and anti-GFP (green), respectively, showing the colocalization of Cx34.1 and Cx34.7 a some GJs (projection of 13-16 sections at 0.71 µm z-step size). (D) colocalization of Cx34.1 and Cx34.7 a some GJs. Insets 1 and 2: GJs containing only Cx34.1 or Cx34.1 and Cx34.7 are interleaved projection of 20 sections at 0.71 µm z-step size). (E) Line scans of fluorescence over distance in two puncta from expanded contact areas showing the presence of Cx34.1 and postsynaptic Cx34.7 (left) or only Cx34.1 (right) at individual GJs. (F) Line scan (fluorescence over distance; normalized to the highest value) of two neighboring puncta in the same expanded contact area showing the presence of Cx34.1 and Cx34.7 in one and only Cx34.1 in the neighboring one. (G,H) Line scans (normalized fluorescence over distance) of GJs showing the presence of both (G), Cx34.1 and Cx34.7 (n=16 from 9 fish) or (H), only labeling for Cx34.1 (n=18 from 9 fish). Thicker lines represent the average of the lines scans for each fluorescence. The scale bars represent actual dimensions; expanded images were not adjusted for expansion factor.

Given the coexistence of the two ZF orthologs of Cx34 at CEs, we asked if the same principle of organization applies to the presynaptic side of the electrical synapse. That is, we asked if both orthologs of Cx35, Cx35.5 and Cx35.1, were present at CEs. For this purpose, we generated a *Cx35.1-mCerulean* transgenic fish, which was double labeled with anti-GFP and anti-Cx35/36, which recognizes both Cx35.5 and Cx35.1 (25). As illustrated in Fig. 5, in contrast to Cx35/Cx36 labeling (Fig. 5B), we didn’t find labeling for GFP (Fig. 5C), indicating the absence of Cx35.1 at CE contact areas. [Traces of GFP labeling were, however, observed in ∼9.1% of the screened lateral dendrites (see Supplemental Fig. 2).] In the absence of significant GFP labeling, the Cx35/36 labeling observed in *Cx35.1-mCerulean* fish reflects the presence and requisite functional role of Cx35.5 at presynaptic sites of CE electrical synapses, that, in addition to Cx35.1 mutation (Fig. 5D, left), it was confirmed with an anti Cx35.5 specific antibody (Fig. 5D, right; see also (25, 33)). The lack of GFP labeling likely represents the pattern of distribution of Cx35.1 in the hindbrain of ZF because: i) the distribution was observed in multiple lines of this transgenic fish, and ii) anti-GFP labeling in Cx35.1-mCerulean fish retina (Fig. 5E) revealed the expected distribution of this Cx in the outer plexiform layer (OPL, Fig. 5F) and inner plexiform layer (IPL, Fig. 5G) of this structure (13).

**Figure 5.**
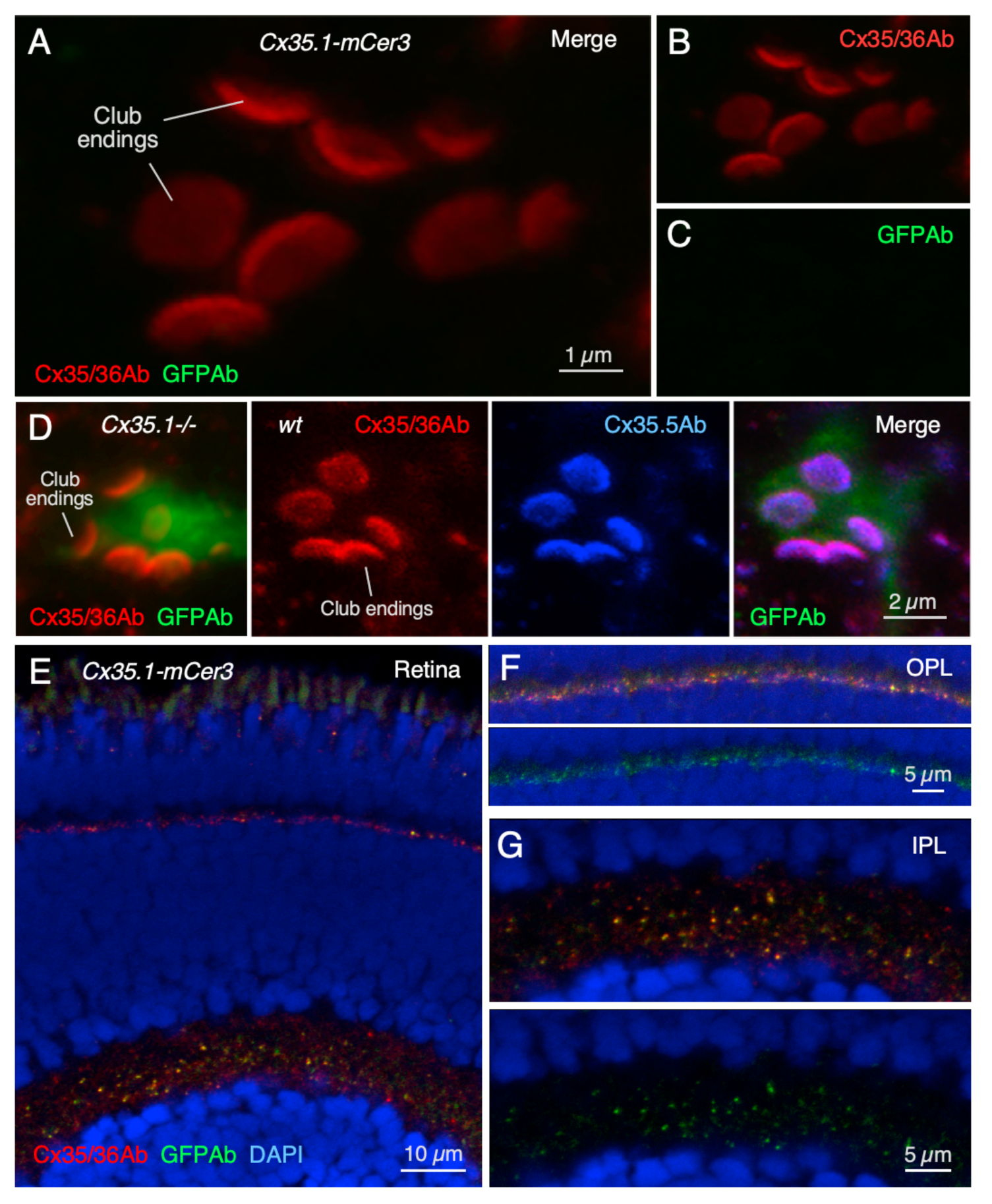
Distribution of Cx35.1 in transgenic fish. (A) CE synaptic contact areas labeled with the Cx35/36 antibody (green), which recognizes both Cx35.5 and Cx35.1, and anti-GFP (green) on a stretch of the lateral dendrite of a Cx35.1-mCerulean (Cx35.1-mCer3) 5 dpf ZF (projection of 10 sections at 0.40 µm z-step size). (B) Club ending contact areas labeled with anti-Cx35/36. (C) Labeling with anti-GFP. (D) Left: Labeling with anti-Cx35/36 (red) and anti-GFP (green, M-cell) in a *gjd2b/Cx35.1-/-* fish (projection of 20 sections at 0.19 µm z-step size). Right: Labeling with anti-Cx35/36 (red) and anti-Cx35.5 (blue) in a *wt (Tol2)* fish, which expresses GFP in the M-cell (green; projection of 12 sections at 0.37 µm z-step size). (E) Labeling with anti-GFP (green) and anti-Cx35/36 (red) reveals the presence of Cx35.1 in the retina of 5 dpf ZF. (F) Distribution of anti-Cx35/36 (top) and anti-GFP (bottom) in the outer plexiform layer (OPL). (G) Distribution of anti-Cx35/36 (top) and anti-GFP (bottom) in the inner plexiform layer (IPL). Labeling was counterstained with DAPI (blue).

Results from the transgenic fish generated in this study unequivocally confirm the presence of Cx35.5 presynaptically at CE contacts and in addition exposed the coexistence of Cx34.1 and Cx34.7 in a subset of GJs, suggesting a functional association between the two postsynaptic Cxs. Surprisingly, despite its presence at CEs, Cx34.7 cannot compensate for the lack of Cx34.1 as shown using *gjd1a/Cx34.1-/-* fish (25). One explanation could be that, despite previous evidence using the perch sequence (39, 40), zebrafish (zf) Cx34.7 is incapable of forming heterotypic channels with zf Cx35.5. To explore this possibility we tested the ability of zf Cx34.1 and zf Cx34.7 to form channels with zf Cx35.5 using exogenous expression in *Xenopus* oocytes and direct measurements of junctional conductance, g_j_, using the dual, two-electrode voltage clamp technique. Both zf Cx34.1 and zf Cx34.7 formed functional homotypic channels and heterotypic channels with zf Cx35.5 (Fig. 6). Homotypic Cx34.1 and Cx34.7 channels differed substantially in their voltage-dependence. Shown are plots of g_j_ vs V_j_, the transjunctional or transsynaptic voltage, obtained by stepping the voltage in one cell while maintaining the voltage in the other cell constant (Fig. 6 A,B). Plotted are normalized, average initial g_j_ and steady-state g_j_ values; the latter represents channel gating whereas initial g_j_ reflects open channel properties. Cx34.1 GJs show little in the way of gating over a range of ±100 mV V_j_ range. In contrast, Cx34.7 exhibits gating with increasing V_j_s with g_j_ declining symmetrically about V_j_ =0. Initial g_j_ shows a modest symmetric increase, similar to that shown for the mammalian neuronal counterpart, Cx36 (41). When the postsynaptic Cxs were paired heterotypically with Cx35.5, the resulting channels showed differential asymmetries exposing a functional difference between GJs hemichannels formed by Cx34.1 or Cx34.7 and Cx35.5. In all the plots, positive V_j_s represent relatively positive voltages on the Cx35.5 side. Heterotypic GJs with Cx34.1exhibit a strong initial (fast) rectification that increases when the Cx35.5 side is made more positive (Fig. 6D). In addition, substantive gating is only observed for one polarity, again positive on the Cx35.5 side. This contrasts the behavior of the Cx34.7/Cx35.5 GJ, which shows modest asymmetries, but with substantive gating occurring for both polarities of V_j_ (Fig. 6C). Thus, the inability of Cx34.7 to compensate for the lack of Cx34.1 at CE contacts is not due to an intrinsic inability to form channels with Cx35.5. Importantly, the different biophysical properties of Cx34.1 and Cx34.7 channels indicate that the presence or absence of Cx34.7 imparts GJs with different functional properties.

**Figure 6.**
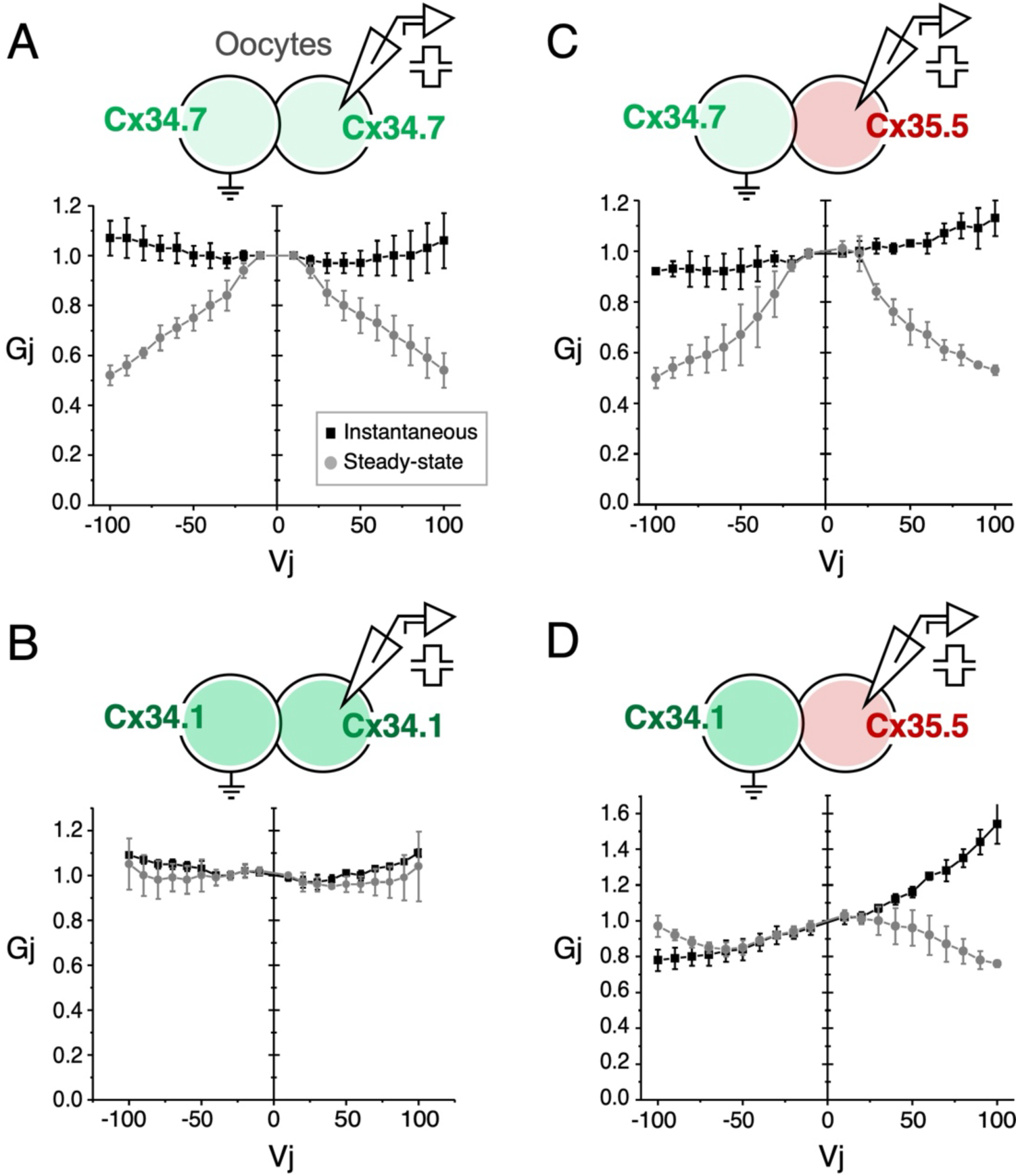
Both Cx34.1 and Cx34.7 can form channels with Cx35.5 when exogenously expressed. Examples of G_j_-V_j_ relationships for homotypic (A,B) Cx34.7 (light green) and Cx34.1(dark green) junctions and heterotypic junctions of each Cx with Cx35.5 (red) (C,D). Plotted are initial (black squares) and steady-state (gray circles) normalized conductances obtained using a dual two-electrode voltage clamp. Voltage steps were applied to the cell indicated. The g_j_ values computed for each cell pair were normalized to values near V_j_=0 obtained by averaging g_j_ values from small V_j_ steps of either polarity. Cx34.1 essentially lacks gating (B) and imparts a strong initial (fast) rectification when docked with Cx35.5 (D). Plotted values represent mean ± SD obtained from several cell pairs (Cx34.1 (n=3), Cx34.7 (n=3), Cx34.7/Cx35.5 (n=4) and Cx34.1/Cx35.5 (n=4).

## DISCUSSION

A perceived difference between chemical and electrical synapses is their molecular and structural complexity, in particular the asymmetry in the organization of the pre- and postsynaptic compartments. Because electrical transmission, unlike chemical transmission, is mediated by intercellular channels formed by corresponding elements, namely Cx hemichannels, on either side of the synapse, a high degree of structural and functional symmetry in their organization has been presumed. Here, we provide evidence of a more complex and asymmetric organization on pre- and postsynaptic sites of a vertebrate electrical synapse formed by two heterologous cell types: auditory afferents that synapse as single contacts known as CEs and the M-cell. Previous ultrastructural analyses indicated different homologs of Cx36 were present at pre vs. postsynaptic sites of this electrical synapse (8, 31), a finding that was independently verified by subsequent genetic studies (24, 25). Our findings here indicate that the asymmetry at this synapse extends beyond the Cx isoforms expressed on either side of the junction, but also the number of isoforms, their configuration at the postsynaptic hemiplaque (Fig.7). Recent structural analysis indicated that the electrical synapse at CE in larval zebrafish is formed by multiple (∼35) GJs that, together with interleaving adherens junctions, occupy ∼80% of the contact’s surface (33). The results here show that these GJs appear to co-exist in at least two configurations. That is, while Cx35.5 is exclusive at the presynaptic hemiplaque, we find that although expression of Cx34.1 is widespread and present in all the GJs at individual CEs, Cx34.7 is co-expressed with Cx34.1 in a subset of GJs.

**Figure 7.**
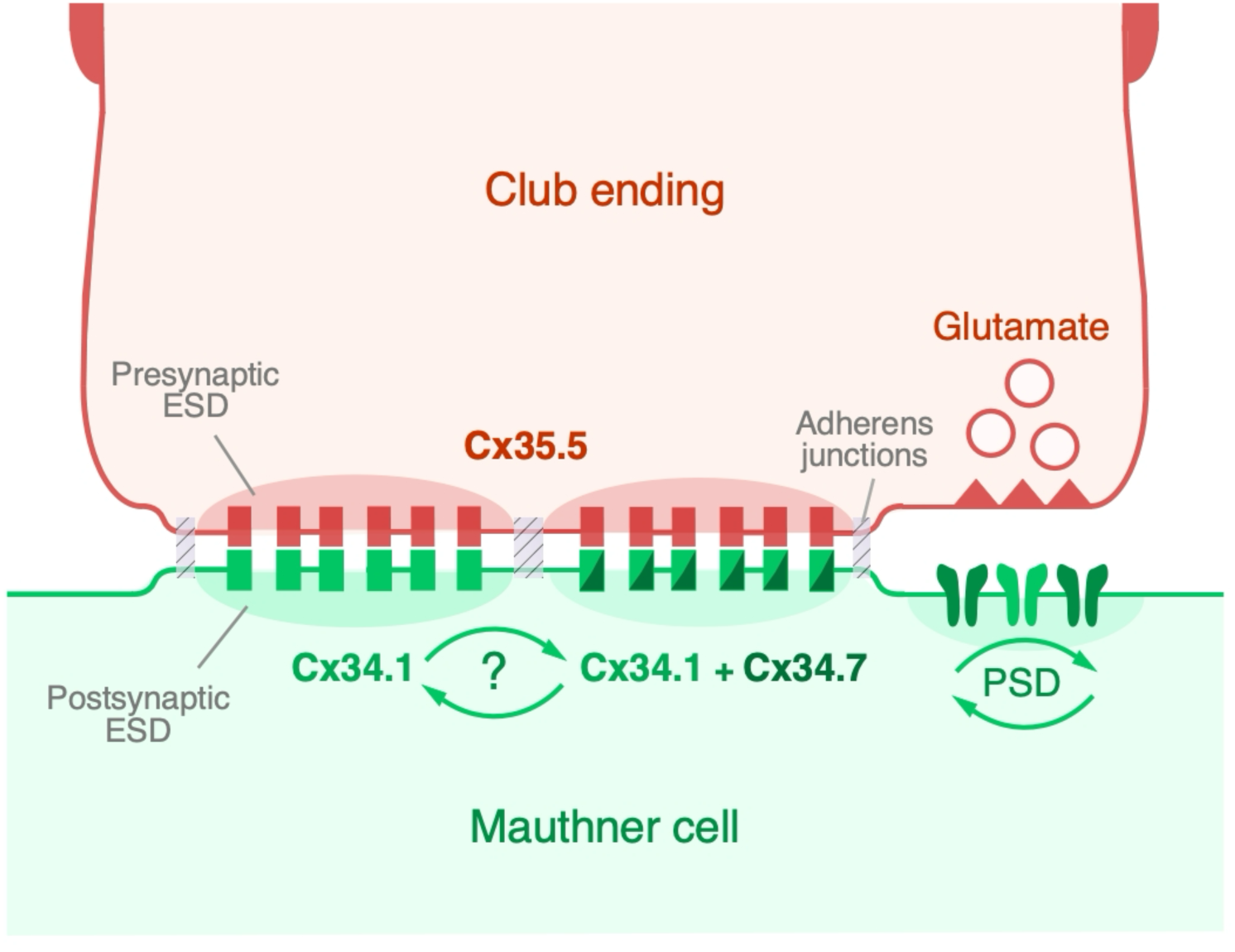
The cartoon summarizes the composition of GJs found at electrical synapses between CE and the M-cell. GJs between CEs (‘Club ending’, red) and the M-cell (‘Mauthner cell’, green) are formed by heterotypic hemichannels and supported by a molecular scaffold (‘Presynaptic ESD’ and ‘Postsynaptic ESD’). Gap junctions are interleaved with Adherens Junctions; see reference (33). The presynaptic CE contributes presynaptic hemichannels made of Cx35.5. In contrast, the M-cell contributes two types of postsynaptic hemichannels. While GJs are formed by hemichannels made of presynaptic Cx35.5 (and postsynaptic Cx34.1 others are formed by hemichannels made of presynaptic Cx35.5 and postsynaptic Cx34.1 (green) and Cx34.7 (dark green). Cx34.1 is obligatory for the formation of the hemichannel and Cx34.7 seems to play an accessory role, likely assembling with Cx34.1 in the postsynaptic hemichannel under specific functional contexts (question mark and arrows). The composition of the postsynaptic GJ hemiplaque, with multiple Cxs, is reminiscent of the organization at chemical synapses at which the PSD regulates receptor types and their relative numbers (arrows) to generate functional diversity.

Asymmetry in the organization of the intercellular GJ channel was shown to influence its functional properties (42). Our biophysical studies here show that Cx34.1 and Cx34.7 exhibit different biophysical properties and impart different behaviors to heterotypic channels formed with Cx35.5, suggesting that the Cx35.5/Cx34.1 and Cx35.5/Cx34.1+Cx34.7 configurations represent two distinct functional states of GJs at CEs. That is, we show here that Cx35.5/Cx34.1 channels express strong rectifying properties (ability to conduct ions more readily in one direction than the other) when compared to Cx35.5/Cx34.7 channels. Thus, the addition of Cx34.7 to individual GJs is likely to modify its properties, either as side-by-side heterotypic channels or combined with Cx34.1 in a heteromeric hemichannel configuration. Rectification of synaptic transmission is a known consequence of heterotypic channels (42–44); the predominance of GJs in the Cx34.1/Cx35.5 configuration suggests that, as reported for goldfish CEs (8), electrical transmission at zf CE exhibits rectification. The properties of Cx34.1/Cx35.5 channels is reminiscent of the rectifying properties of single CE contacts in goldfish (8) indicating that, in addition to Cx34.7, Cx34.1 might also be present at CEs of this species. [The existence of two orthologs of Cx34 in teleost fish was then unknown and more recently exposed by genetic analysis in zf (30).]

We note that other channel properties, in addition to rectification, may differ between Cx34.1 and Cx34.7. These include unitary conductance and open channel lifetimes that impact the strength of coupling thereby playing a role in modulating the efficacy of electrical transmission. Given the importance of the first extracellular loop, E1, domain of Cxs in determining channel gating and permeability characteristics (45–48), sequence comparison of zf Cx34.1 and Cx34.7 shows an overall high conservation (83% identity, 91% similarity), but a notable difference at position 44 in the E1 domain; K44 vs T44 in Cx34.1 and Cx34.7 respectively. This position lies close to D47, the putative site for the binding of Mg^2+^, which has been shown to be a potent regulator of channel conductance unique to Cx36 among Cxs and verified in perch orthologs, Cx35 and Cx34.7 (49). Incorporating Cx34.7 subunits in homomeric or heteromeric configurations in postsynaptic hemichannels could affect Mg^2+^ binding, offering a mechanism of plasticity in addition to modulation of channel numbers (50). Expression of Cx34.7 could also offer other means of regulation through changes in binding with scaffolding proteins, kinases and other modulatory factors. Given the similarities in overall sequence, it is likely that Cx34.1 and Cx34.7 assemble into heteromeric hemichannels with varying subunit combinations. Intermingled channels consisting of homomeric postsynaptic hemichannel assemblies would also impart changes in transmission. Thus, the inclusion of Cx34.7 in GJs likely results in changes in the functional state of the CE electrical synapse. Finally, the coexistence of Cx34.7 and Cx34.1 does not appear to be developmentally regulated as: i) Cx34.7 and Cx34.1 are not functionally redundant and interchangeable, given the different biophysical properties that we describe here, and ii) the structural and functional properties of CE electrical synapses were found conserved between adult goldfish and zebrafish larvae (8, 35, 37, 51).

### A novel connexin functional arrangement at electrical synapses

Remarkably, despite its presence, our previous data shows that Cx34.7 cannot compensate for the loss of transmission in *gjd1a/Cx34.1-/-*fish, whereas transmission is unperturbed in *gjd1b/Cx34.7-/-* fish (25). This finding reveals a novel functional configuration for Cxs at neuronal GJs, in which while the presence of Cx34.1 is obligatory, Cx34.7 seems to serve a non-obligatory, regulatory, role. Consistent with a regulatory function, a likely role for Cx34.7 would be to form channels with Cx34.1 in a heteromeric configuration, presumably in certain functional contexts, to modify channel properties. Although such a Cx functional configuration could result from the inability of zf Cx34.7 to form channels with zf Cx35.5, our data indicate that both zf Cx34.1 and zf Cx34.7, when exogenously expressed, can form heterotypic channels paired with zf Cx35.5. The functional role of Cx34.7 at CE electrical synapses is therefore determined by mechanisms other than the intrinsic ability of this Cx for form channels with Cx35.5. The functional assignment could be likely specified by the postsynaptic regulatory molecular machinery. This possibility is consistent with recent evidence indicating the molecular complexity and functional hierarchy of the molecular scaffold over Cx channels at neuronal GJs (7, 9, 25, 52, 53). For example, the scaffolding protein zonula occludens-1 (ZO1) was shown to be critically required for the presence of Cxs and synaptic transmission at CE synapses, whereas in contrast, Cxs were not required for the presence of ZO1 at these contacts (25). While the molecules responsible for the functional assignment of Cx34.7 are currently unknown, recent proteomic analyses obtained by proximity-dependent biotinylation linked to neural Cxs in zebrafish and mice identified hundreds of additional Cx-associated proteins (12, 13), some of which are likely to participate in the support and plastic regulation of electrical transmission. These proteins are likely to be part of the electrical synapse density (ESD; (25)) (Fig.7), a molecular structure comparable the postsynaptic density (PSD) of chemical synapses, characteristically observed in EM images of neuronal GJs (54).

### Electrical and chemical synapses share a common postsynaptic motif

The GJs at CE electrical synapses are between two different neurons and, therefore, inherently asymmetric (Fig. 7). Structurally, the GJ ‘plaque’ (55) consists of the apposition of two ‘hemiplaques’, containing the pre- or the postsynaptic hemichannels that together form the communicating intercellular GJ channels. Interestingly, the presynaptic CE terminal and postsynaptic M-cell dendrite seem to follow different strategies of organization at their GJ hemiplaques. That is, the structural complexity of postsynaptic organization contrasts with the apparent simplicity of the presynaptic GJ hemiplaque with a main single Cx forming the presynaptic hemichannel (Fig. 7). The composition of the postsynaptic GJ hemiplaque, operating with multiple Cxs, is reminiscent of the postsynaptic organization of chemical synapses, at which the PSD regulates the receptor types and their relative numbers to generate functional diversity (14, 56). Emphasizing the asymmetric organization of the postsynaptic hemiplaque, the ESD scaffold ZO1 was found to be exclusively located postsynaptically, where it functions to regulate the presence of Cxs (25, 33), and Neurobeachin was shown to function postsynaptically, contributing to the delivery and specificity of Cx localization (11, 57).

The shared organizational motif between postsynaptic GJ hemiplaques and PSDs (Fig. 7) is supported by the functional relationship between these two structures at CEs, during activity-dependent changes in synaptic strength (19, 21, 58). Recent analysis identified overlapping proteins between the PSD and the Cx proteomes (12), suggesting that a structural association might support the functional relationship between glutamatergic PSDs and the postsynaptic hemiplaque of neuronal GJs. Interestingly, the relative proportion of GJs with coexisting Cx34.1 and Cx34.7 is variable between neighboring CE contacts, suggesting this GJ configuration could be dynamically regulated. The variable expression and conditional nature of Cx34.7 is reminiscent of the distribution of CaMKII observed at goldfish CEs (59), where it associates with Cxs at individual GJs (59). Electrical (and chemical) transmission at CEs undergoes activity-dependent potentiation mediated by postsynaptic activation of CaMKII at PSDs nearby GJs (21, 58), and the presence and amount of CaMKII was proposed to be indicative of synaptic strength, which is also variable between individual CEs (58, 59). While it remains speculative, the presence of Cx34.7 could be associated with changes in synaptic strength known to take place at these terminals. Thus, the overall functional state of the CE electrical synapse could reflect the sum of GJs co-existing in each functional configuration (Fig.7).

The large size and colocalization with glutamatergic transmission at mixed contacts, less common in mammals (although see (60)), might make CEs to be perceived as atypical electrical synapses. However, functional features first identified at these contacts were reported to occur at electrical synapses elsewhere (27, 28, 61–64). The genes *gjd1* (Cx34) and *gjd2* (Cx35) can be found in most vertebrate species including fish, birds and amphibia (30), forming in most cases heterotypic junctions (65). However, mammals only have one gene, *gjd2*, which encodes for Cx36, since *gjd1* (Cx34) was absent from the mammalian genome. Cx36 is widely distributed in the mammalian brain (66, 67) and responsible for transmission at most electrical synapses (68). The fact that mammalian electrical synapses exhibit biophysical properties and plastic mechanisms (28, 61, 62, 69) parallel to those observed in fish CEs (27–29, 62), suggests the existence of molecular mechanisms at the RNA or protein level capable of creating a similar functional diversity functioning only with the *gjd2* gene. Thus, the properties and combinations of fish Cx36 homologs could therefore serve to expose a hidden diversity in the organization of electrical transmission elsewhere.

## METHODS

All experiments were performed in 5 days post fertilization (dpf) zebrafish (ZF), *Danio rerio*. The fish were housed at the zebrafish facility in the Department of Neuroscience of the Albert Einstein College of Medicine, and bred and maintained at 28°C on a 14-hour light / 10-hour dark cycle.

### Immunohistochemistry

ZF larvae were anesthetized with 0.03% MS-222 (tricaine methanesulphonate) and fixed for 3 hours in 2% trichloroacetic acid, in 1X PBS, at room temperature. Fixed samples were then washed 3 times with 1X PBS. The brain was the dissected with the help of a custom-made tungsten needle and forceps. This needle was also used to remove the cerebellum, optic tectum, and telencephalon with the purpose of improving the working distance of the microscope objective (unnecessary in samples used for expansion methods). The dissected brains were washed with 1X PBS + 0.5% Triton X-100 (PBS-Trx) and blocked with 10% Normal Goat Serum + 1% DMSO in PBS-Trx. Brains were then incubated overnight with a primary antibody mix and kept at room temperature. This antibody mix (which included block solution), contained combinations of the following: rabbit anti-Cx35.5 (Miller et al., 2015 (44), clone 12H5, 1:200), mouse IgG1 anti-Cx35/36, which labels both Cx35.5 and Cx35.1 (Millipore, cat. MAB3045, 1:250), mouse IgG2A anti-Cx34.1 (Miller et al., 2015 (44), clone 5C10A, 1:200), and chicken IgY anti-GFP (Abcam, cat. ab13970, 1:200), and rabbit IgG anti-GFP (Invitrogen, cat G10362, 1:200). Brains were then washed 3 times in PBS-Trx, and then blocked with 10% Normal Goat Serum + 1% DMSO in PBS-Trx, followed by incubation in a secondary antibody mix at room temperature for 4 hours. The secondary antibody mixes included, in addition to block solution, combinations of antibodies raised in goat against mouse IgG- Alexa Fluor 546 (Invitrogen, cat. A11030, 1:200), mouse IgG- Alexa Fluor 647 (Invitrogen, cat. A21235, 1:200), rabbit IgG- Alexa Fluor 546 (Invitrogen, cat. A11010, 1:200), rabbit IgG- Atto 647N (Sigma-Aldrich, cat. 40839, 1:200), mouse IgG- Atto 647N (Sigma-Aldrich, cat. 50185, 1:200), and chicken IgY-Alexa Fluor 488 (Invitrogen, cat. A11039, 1:200), rabbit IgG-Alexa Fluor 488 (Invitrogen, cat. A11008, 1:200). Samples were washed with 1X PBS 4 times and overnight at 4°C, and then transferred in the dark, onto a slide, and mounted with either Fluoromount-G (Invitrogen, cat.00-4958-02) or ProLong Gold antifade (Invitrogen, cat. P36930). Samples were then covered using the ‘bridge’ procedure (67) and sealed with nail polish. In the case of the samples used in expansion microscopy, expansion procedure the samples were instead incubated in an anchoring solution (see below) overnight.

### Expansion procedure

Expansion of brain samples was performed following previous protocols (70) with additional modifications as reported in Cardenas et al., 2024 (22) and Ijaz et al., 2024 (38). The following reagents were used (final concentrations reported): Anchoring solution: Acryloyl-X, SE (0.01%) and PBS (1X) to a complete volume. Monomer solution/gelling solution (4-HT, TEMED, and APS should be added sequentially, one at a time, to each sample): acrylamide (2.5%), N,N’ methylenebisacrylamide (0.15%), NaCl (2M), PBS (1X), Sodium Acrylate (8.6%), 4-hydroxy-TEMPO (4-HT) (0.01%), Tetramethylethylenediamine (TEMED) (0.2%), Ammonium Persulfate (APS) (0.2%), and cell culture grade water to a complete volume. Digestion buffer: Tris (pH 8.0) (50 mM), EDTA (1 mM), Triton X-100 (0.5%), NaCl (0.5 M), and cell culture grade water to a complete volume, and Proteinase K (8 units/mL). Following the above described immunolabeling steps, the sample was incubated in anchoring solution overnight. After this, the sample was washed twice in 1X PBS and placed into monomer/gelling solution, first at 4°C for 50 minutes, and then at 37°C for 2 hours. Once the hydrogel polymerized, the sample was placed at 50°C for 12 hours in the digestion buffer + Proteinase K. Following this procedure, the samples were then washed 5 times, each being ten minutes on a shaker plate, with cell culture grade water for expansion and imaged with the help of the confocal microscope.

### Confocal imaging

All images were acquired with a LSM 710 and a LSM 880 Zeiss microscopes, using the following laser wavelengths: Argon 458/488/514, HeNe 543, and HeNe 633, along with the corresponding filter: MBS 488/543/633, MBS 458/543, MBS 488/543/633, using either a 40×1.0 NA water immersion objective or a 63x 1.40 NA oil immersion objective. Both lateral and ‘En face’ views of expanded CE contact areas were used for quantitative analysis. Confocal settings were adjusted to achieve maximum visualization of labeling at CE contact areas. The lateral and axial resolutions of our LSM 710 system were estimated following the microscope specifications (71) to range between 248.9 to 322 nm and 429.44 nm to 557.04 nm, respectively, depending on the excitation wavelength used (488, 543, 633). Following corrections for the linear expansion factor (3.9x), the lateral and axial resolutions after expansion are expected to range between 63.81 nm to 82.77 nm and 110.11 nm to 142.83 nm, respectively.

### Quantification and statistical analysis

Confocal images were obtained using ZEN (black edition) software and analyzed using FIJI. Expansion images in the figures were not adjusted for expansion factor and therefore the scale bars represent actual dimensions. The contrast and brightness of the fluorescence channels were individually adjusted and contrasted in Photoshop (Adobe), including the blur and sharpen filters. However, quantitative analysis of image fluorescence for double labeling analysis was carried out in the raw data (see below).

### Colocalization analysis

The analysis of labeling colocalization was performed in Fiji (https://imagej.net/imaging/colocalization-analysis) using the JACoP plugin (https://imagej.net/plugins/jacop, which uses pixel-wise methodology to determine matching pixels between channels. Following selection of the ROIs of oval CE contact areas, colocalization was quantified using the Manders’ coefficient (72, 73). The Mander’s coefficient expresses the proportion of fluorescence in one channel that colocalizes with the proportion of fluorescence in the second channel, and vice-versa (74). The procedure allows setting independent thresholds for each channel to account for different levels of fluorescent intensity to then generate colocalization coefficients whose values range from 0 to 1.

### Retina immunohistochemistry

Whole eyes were removed from euthanized zebrafish and fixed in 4% PFA in 0.1M phosphate buffer, pH 7.5 for 30 minutes at room temperature. Afterwards, the eyes were washed three times for 10 minutes each and cryoprotected overnight in 30% sucrose in PBS. On the next day, whole eyes were embedded in Tissue-Tek O.C.T. cryomatrix (Sakura Finetek, Torrance, CA, 530 USA) and stored at -20°C. Afterwards, whole eyes were cut into 20µm thick cryosections and incubated at 37°C. Each section was incubated overnight at 4°C in the primary solution containing 10% normal donkey serum in 0.5% Triton X-100 in PBS. The following primary antibodies were used: Cx35, 1:250 Clone: 8F6, #MAB3045, Millipore and GFP, 1:250, #A-11122, Thermofisher. On the next day, retinal sections were washed three times for 10 minutes and incubated for 1h at RT in the secondary antibody solution containing 10% normal donkey serum in 0.5% Triton X-100 in PBS. The following secondary antibodies were used: Donkey anti-mouse conjugated with Cy3, 1:250, #715-165-150, Jackson ImmunoResearch and donkey anti-rabbit conjugated with Alexa488, 1:250, #711-545-152, Jackson ImmunoResearch. After the incubation, sections were washed three times for 10 minutes each and mounted with Vectashield containing DAPI, H-2000, Vector Laboratories Inc.

### Imaging of retina sections

Images of retina sections were acquired with a confocal laser scanning microscope (Zeiss LSM 800) using a 60x oil objective. Image dimensions: 1024×1024 pixels, pixel size: 66nmx66nm. Confocal scans were processed using Fiji (ImageJ2 version 2.14.0/1.54f) software and slightly adjusted in brightness and contrast.

### Electrophysiology

Synaptic responses were obtained during whole-cell recordings of M-cells in wt and transgenic zebrafish larvae (5–7 dpf). Details of this procedure can be found in Echeverry et al., 2022 (35). Briefly, fish were first anesthetized with a 0.03% solution of MS222 (pH adjusted to 7.4 with NaHCO_3_) and then transferred to external solution containing d-tubocurarine (10 μM, Sigma). The external solution contained (in mM): 134 NaCl, 2.9 KCl, 2.1 CaCl_2_, 1.2 MgCl_2_, 10 HEPES, 10 Glucose, pH adjusted to 7.8 with NaOH (31). Zebrafish larvae were placed on their backs onto a Sylgard-coated small culture dish (FluoroDish, WPI) and the hindbrain was exposed ventrally. The larva was then placed on an Axio Examiner upright microscope (Carl Zeiss AG) equipped with a recording set-up and superfused with external solution during the entire recording session. The M-cells were identified by GFP expression and/or far-red DIC optics. Patch pipettes (3-4 MΩ) were filled with internal solution containing (in mM): 105 K-Methanesulfonate, 10 HEPES, 10 EGTA, 2 MgCl_2_, 2 CaCl_2_, 4 Na_2_ATP, 0.4 Tris-GTP, 10 K_2_-Phosphocreatine, 25 mannitol, pH adjusted to 7.2 with KOH. Whole-cell recordings were performed under the current-clamp configuration with the help of a Multiclamp 700B amplifier and a Digidata 1440A (Molecular Devices) digitizer. The liquid-liquid junction potential (estimated in -16 mV using Clampe× 10.6; Molecular Devices) was subtracted from the measured values. For extracellular stimulation of the auditory afferents terminating as CEs on the M-cell, a septated (theta) glass pipette was filled with external solution and placed near the posterior macula of the larva’s ear, where the cell bodies of the of auditory afferents cluster together (11). The maximal amplitude of the electrical and chemical components was estimated by applying shocks of increasing intensity until the amplitude of the electrical component did not further increase and before additional responses with longer latency were evoked. For estimates of the M-cell input resistance, a hyperpolarizing-current step of -1 nA and 20 ms in duration was intracellularly applied through the recording electrode and the evoked voltage deflection measured, followed by derivation of input resistance by Ohm’s law.

### Exogenous expression of connexins

All zebrafish Cx constructs were generated using a gene synthesis service (GenScript, Piscataway, NJ) and cloned into the BamHI restriction site of the pCS2+ expression vector for functional studies and exogenous expression. All constructs were verified by sequencing. For expression in *Xenopus laevis* oocytes, RNA was synthesized and oocytes were prepared and injected as previously described (45, 75). RNA concentrations were measured using a NanoDrop 2000 Spectrophotometer (Thermo Scientific, Waltham MA) and tubes containing equal concentrations were prepared by adding appropriate amounts of nuclease free water. 50 nL volumes of RNA were injected using a Nanoject II auto-nanoliter injector (Drummond Scientific Company, Bromall PA). Injected oocytes were maintained at 16 – 18 °C in a modified ND96 solution (MND96) containing (in mM) 100 NaCl, 2 KCl, 1 MgCl_2_, 1.8 CaCl_2_, 10 glucose, 10 HEPES, 5 pyruvate, pH adjusted to 7.6. *Xenopus laevis* oocytes were purchased from Xenopus 1 (Dexter, Michigan).

### Electrophysiological recordings

To measure gap junctional conductance (g_j_) *Xenopus* oocytes were manually devitellinized and paired by placing them in dishes coated with agarose (1%). Each oocyte of a pair was clamped independently using a two-electrode voltage clamp to a common potential. Transjunctional voltage, V_j_, steps were applied by stepping the voltage in one cell of a pair; the voltage in the other cell remained constant. Voltage protocols to obtain gj-Vj relationships consisted of voltage steps to one cell, 10 s in duration, ranging over ± 100 mV in 10 mV increments. Each V_j_ step was preceded by a small, brief pre-pulse of constant amplitude so that a family of currents could be normalized if the expression level changed over the course of an experiment. Currents were filtered at 200 Hz and digitized at 1 -2 kHz. Steady-state g_j_-V_j_ relationships were obtained by extrapolating exponential fits of the data from voltage steps to t = ∞ as previously described (44). Only cell pairs with g_j_ values ≤ 10 µS were used in generating g_j_-V_j_ relationships to avoid effects of series access resistance on voltage dependence (76).

### Connexin clones

Expression and transgene clones for each of the zebrafish (ZF) connexins were generated from genomic bac or fosmid clones. Cx35.1 clones were derived from chromosome 20 bac RP71-1C22; Cx35.5 clones were derived from chromosome 17 fosmid ZFishFos1141d03; Cx34.1 clones were derived from chromosome 5 bac CH211-87N9; Cx34.7 clones were derived from chromosome 15 bacs CH1073-29P7 containing exon 1 and CH73-388K15 containing exon 2. For each connexin, a pseudo-cDNA clone was generated by PCR amplification of exons 1 and 2 and assembly into a cloning vector using conventional cloning or Cold Fusion (System Biosciences, Palo Alto, CA). Cx35.1 was cloned first into transcription vector SP64T and then transferred into the vector backbone derived from mammalian expression vector pEGFP-N1 by deletion of EGFP with AgeI and NotI restriction enzymes. The other three connexins were cloned directly into the N1 vector backbone creating untagged mammalian cell expression clones.

To create visible connexins, we tagged each of the ZF Cxs with a fluorescent protein. Critically important in this design, the fluorescent protein coding regions were inserted into the CT of the connexin genes at a site (19-21 aa before end of the CT) that was previously found not to disrupt regulation of coupling by connexin phosphorylation and not to block the C-terminal PDZ-domain interaction motif (36). Cx35.1 and Cx35.5 (presynaptic Cxs) were tagged with mCerulean3; Cx34.1 and Cx34.7 (both postsynaptic Cxs) were tagged with oxGFP. All clones were tested for proper folding, trafficking and gap junction formation by transfection into Hela cells (CCL2; ATCC, Manassas, VA).

Transgene clones were created in a Tol2 transposon-based vector (77) that has been modified to include a cassette containing mCherry driven by a 250 bp *myl7* promoter to produce a visible red label in the heart (78). For each clone, the FP-tagged pseudo-cDNA was cloned into the Tol2 vector and genomic elements upstream of the connexin open reading frame added. The Cx35.1 construct contained 4.6 kb of sequence upstream of exon 1 plus the full exon 1 and intron 1 cloned into the native BamHI site at the beginning of exon 2. The Cx35.5 construct contained 5.1 kb of sequence upstream of exon 1. The Cx34.1 construct contained 1.9 kb of sequence upstream of exon 1. The Cx34.7 construct contained 2.9 kb of sequence upstream of exon 1 plus a rabbit b-globin intron derived from the Tol2 vector. All promoter fragments were amplified by PCR using Phusion DNA polymerase (New England Biolabs, Ipswich, MA) and cloned using Gibson Assembly (NEB).

### Zebrafish

Stable lines of ZF in which the GJ forming proteins responsible for mediating electrical transmission at mixed synapses on the M-cells: Cx35.1 (AE 39, AE54 and AE5855), Cx34.1 (AE 38, AE35 and AE37) and Cx34.7 (AE5857and AE5856), are tagged with fluorescent proteins were generated in the Albert Einstein Zebrafish Facility using the Tol2 transposon system. Transgenic ZF were made in AB background (ZL1; Zebrafish International Resource Center, Eugene, OR). All the transgenic ZF were generated and bred at Zebrafish Core of the Albert Einstein College of Medicine. Two other previously described lines were also used: *Tol056* transgenic line, which carry a GFP fluorescent marker under control of the promoter for the Heat Shock Protein 70, and, which include the M-cells (79), and Cx35.1-/- line (*gjd2b(fh454*) containing a gjd2b^delta12bp deletion (https://zfin.org/ZDB-ALT-170822-2#’variants’).

## ACKNOWLEDGMENTS

We thank Cheryl K. Mitchell for assistance testing expression of connexin clones and Shunichi Yoshikawa for testing expression of zf connexin promoters, and members of the Pereda lab for critical feedback on the work and manuscript. This work was supported by NIH grants R01DC011099 from the National Institute on Deafness and Other Communication Disorders (NIDCD) and R21NS085772 from the National Institute of Neurological Disorders and Stroke (NINDS) to A.E.P., R01EY012857 from the National Eye Institute (NEI) to J.O.B., and RF1MH120016 from the National Institutes of Mental Health (NIMH) to A.E.P. and J.O.B.

## AUTHOR CONTRIBUTIONS

H.H., S.I., V.V. and A.E.P. contributed to experimental design, each with particular emphases. H.H. and S.I. performed and analyzed M-cell immunolabeling and expansion experiments; F.A.E. generated transgenic fish and performed M-cell electrophysiology. J.O.B. and Y-P.L. designed and generated the constructs used in for transgenesis. S.T. performed retina immunolabeling; V.V. performed and analyzed biophysical experiments; A.E.P. oversaw all aspects of the project. All authors wrote the paper and approved of the manuscript.

## DECLARATION OF INTERESTS

The authors declare no competing interests.

---

**Supplemental Figure 1.**
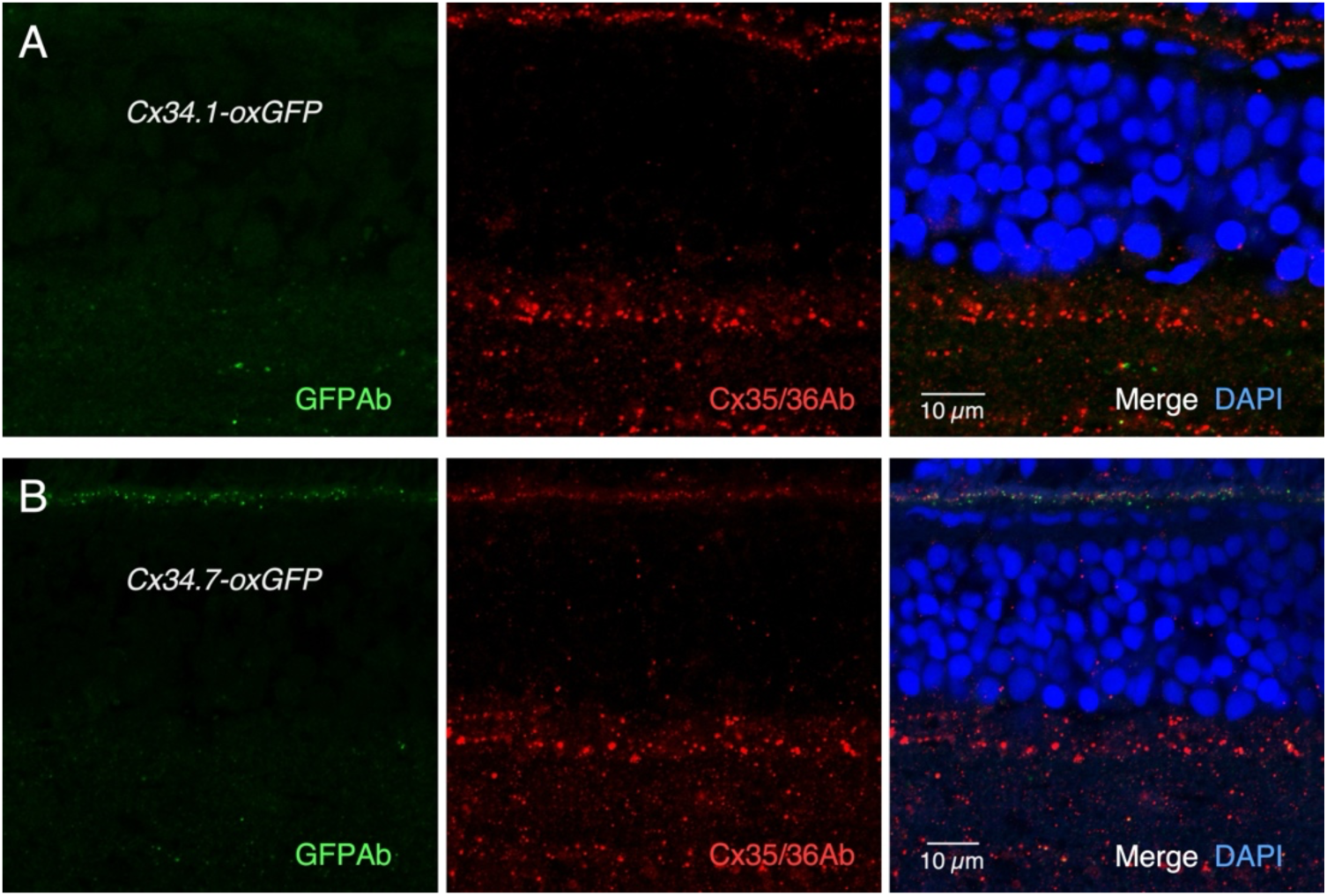
Labeling in Cx34.1-oxGFP and Cx34.7-oxGFP retina. (A) Labeling with antiGFP (left, green) and Cx35/36Ab (center, red) and displayed merged with DAPI counterstain (right, blue) in an adult Cx34.1-oxGFP fish. (B) Labeling with antiGFP (left) and Cx35/36Ab (center) and merged with DAPI counterstain (right) in an adult Cx34.7-oxGFP fish (single confocal sections).

**Supplemental Figure 2.**
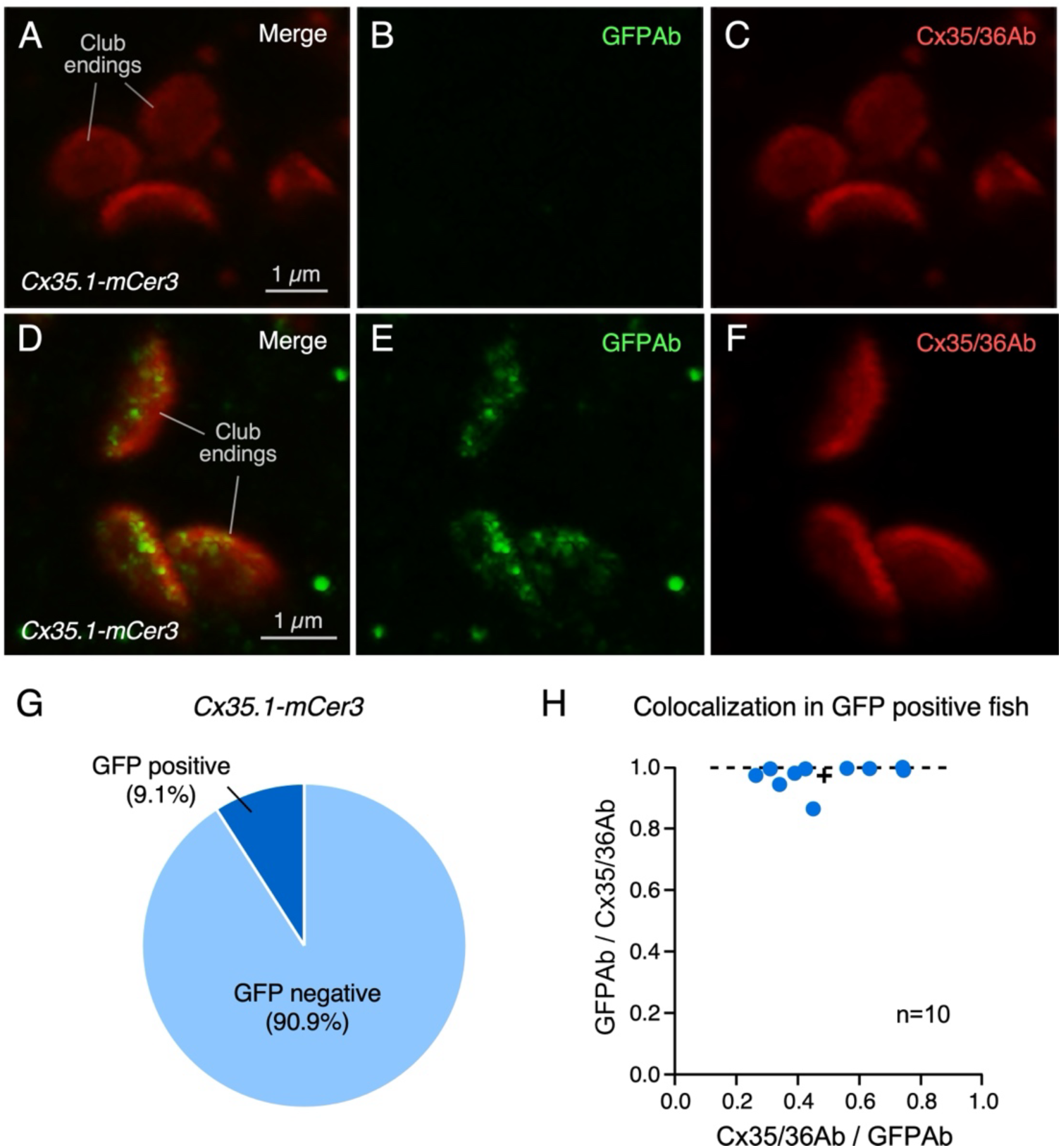
Quantification of labeling in Cx35.1-mCer3 transgenic fish. (A,B,C) About 90% of the Cx35.1-mCer3 fish labeled with anti-Cx35/36 (red) and anti-GFP (green) did not show the presence of Cx35.1 in the M-cell (projection of 10 sections at 0.40 µm z-step size). Some GFP labeling was seen in about 9% of the fish. (B,D,E) shows one of the only examples where significant labeling for GFP was observed at CE synaptic contact areas (projection of 14 sections at 0.40 µm z-step size). (G) Pie plot represents the percentage of M-cell lateral dendrites obtained from three different lines of Cx35.1-mCer3 fish in which traces of GFP labeling was observed (dark blue, ‘GFP positive’, 9.1%). GFP was not detected in 90.9% of the dendrites (n=10 from 6 fish). (H) Colocalization of GFP and Cx35/36 fluorescence at individual CEs Cx35.1-mCer3 fish determined by the Mander’s Coefficient in the examples in which this estimate was possible: GFPAb / Cx35/36Ab 0.973 ± 0.042 (x axis); Cx35/36Ab / GFPAb 0.486 ± 0.175 (y axis), n=10 CEs from 6 fish.

## Notes

### Competing Interest Statement

The authors have declared no competing interest.

